# ADO-MEDIATED SYNTHESIS OF TAURINE ALTERS THE CHROMATIN LANDSCAPE OF INGUINAL ADIPOSE TISSUE TO ENHANCE NON-SHIVERING THERMOGENESIS

**DOI:** 10.1101/2023.02.02.526843

**Authors:** Pei-Yin Tsai, Bo Shui, Seoyeon Lee, Yang Liu, Yue Qu, Chloe Cheng, Kaydine Edwards, Callie Wong, Ryan Meng-Killeen, Paul Soloway, Joeva J Barrow

## Abstract

Non-shivering thermogenesis (NST) has strong potential to combat obesity, however, a safe molecular approach to activate this process has not yet been identified. The sulfur amino acid taurine has the ability to safely activate NST and confer protection against obesity and metabolic disease in both mice and humans, but the mechanism of action is unknown. In this study, we discover that a suite of taurine biosynthetic enzymes, especially that of cysteamine dioxygenase (ADO), significantly increases in response to β_3_ adrenergic signaling in inguinal tissues (IWAT) in order increase intracellular concentrations of taurine. We further show that ADO is critical for thermogenic mitochondrial function as its ablation in thermogenic adipocytes significantly reduces taurine levels which lead to declines in mitochondrial oxygen consumption rates. Finally, we demonstrate via assay for transposase-accessible chromatin with sequencing (ATAC-Seq) that taurine supplementation has the ability to remodel the chromatin landscape to increase the chromatin accessibility and transcription of genes, such as glucose-6-phosphate isomerase 1 (Gpi1), that are critical for NST. Taken together, our studies highlight a potential mechanism for taurine in the activation of NST that can be leveraged toward the treatment of obesity and metabolic disease.

## 1. INTRODUCTION

Obesity is defined as having a body mass index (BMI) greater than or equal to 30 kg/m^2^ and is among the current leading prevalent health issues worldwide. It is well established that obesity is linked with comorbidities such as cardiovascular disease, type 2 diabetes mellitus, and dyslipidemia [1]. According to the Centers for Disease Control and Prevention (CDC) in the United States, the obesity prevalence increased from 30.5% to 41.9% during the period 1999-2020 [2]. Current treatment options, including calorie restriction, bariatric surgery, and pharmacotherapy either present with poor long-term efficacy and/or serious side effects [3-6]. One attractive option to combat obesity is to take advantage of the molecular properties of thermogenic brown and beige adipocytes to raise energy expenditure by activating the non-shivering thermogenesis (NST) program. Brown and beige adipocytes contain specialized mitochondria enriched with the uncoupling protein 1 (UCP1) that is able to convert the dissipation of chemical energy into heat production and maintain the core body temperature [7-9]. In adult humans, thermogenic brown adipose tissue (BAT) is located in the supraclavicular, axillary, and mediastinal regions and studies have shown that when BAT is activated after a 2-hour exposure in a mild cold environment, resting metabolic rates are significantly elevated [10]. Similarly, the ^18^F-fluorodeoxyglucose (^18^FDG) uptake, detected by a positron-emission tomography and computed tomography (PET-CT), was significantly increased in brown adipocytes (BAT) which is a marker of high metabolic activity [11]. While there have been significant advances in the mechanisms that govern the NST activation process such as cold environmental stimuli or the administration of pharmacological agents such as the β_3_ agonist CL 316,243, practical and safe approaches to stimulate NST in humans are still challenging [12,13]. Intriguingly, researchers have discovered that the sulfonic amino acid taurine (2-aminoethanesulfonate) has the natural ability to activate NST to confer protection against obesity in rodent models but the mechanism of how this occurs is unknown [14].

Taurine is a naturally occurring amino acid derived from animal protein or by endogenous synthesis. Unlike the majority of other amino acids, taurine is not a building block of protein synthesis. Instead, it has two major reported roles in humans. The first is its role in bile acid conjugation which can complete the ionization of bile and enhance the emulsification of lipids [15]. In addition, studies have demonstrated that taurine supplementation can stimulate the 7α-hydroxylase (CYP7A1) mRNA expression to increase endogenous bile acid synthesis [16,17]. The other reported role for taurine is the modification of mitochondrial tRNAs where a taurine methyluridine (τm^5^U) becomes incorporated at the wobble position in the anticodon loop of human mitochondrial tRNAs critical for the synthesis of mitochondrial proteins [18,19]. In addition to these defined roles for taurine, the simple sulfur amino acid has been implicated in a host of diverse physiological functions. Indeed, supplementation of taurine is positively correlated with enhanced muscle tone and strength, improved antioxidation, and protection against obesity and metabolic disease via the stimulation of non-shivering thermogenesis (NST) [14,20-22]. In randomized control human trials, researchers discovered a negative correlation between taurine and obesity, and they demonstrated that the supplementation of taurine can significantly decrease the body weight in obese volunteers [23-26]. In mouse studies, the administration of taurine to high-fat diet-induced (HFD) obese mice significantly reduced body weight gain and increased the expression of thermogenic markers, such as uncoupling protein 1 (*Ucp1)* and peroxisome proliferator-activated receptor-gamma coactivator 1 alpha (*Pgc1α)* markers in the inguinal and white adipocytes corresponding to increased oxygen consumption [14,27]. Moreover, taurine supplementation can also decrease the adipogenesis related markers in white adipocytes, such as peroxisome proliferator-activated receptor alpha (PPAR-1α), peroxisome proliferator-activated receptor gamma (PPAR-γ), and CCAAT/enhancer binding proteins alpha (C/EBPα) [28]. The relationship between taurine and NST is reciprocal. Taurine supplementation has the ability to activate NST in mice and correspondingly, activated NST leads to the enhanced synthesis of taurine. Indeed, several studies have demonstrated that pharmacological activation of NST using the β_3_ agonist CL 316,243 can significantly elevate taurine levels in the inguinal and white adipocytes [29]. Overall, although the anti-obesity effect of taurine has been well reported, the underlying molecular mechanism of how this occurs and the association of taurine with NST are still enigmatic.

In this present study, we present a mechanistic role for taurine in the activation of NST. We demonstrate that the taurine biosynthetic enzymes in the inguinal adipose tissue are the most responsive to pharmacological NST activation and that the protein levels of the biosynthetic enzyme cysteamine dioxygenase (Ado) is the most responsive to β_3_ adrenergic stimulation. We further demonstrate that the loss-of-Ado significantly blunts intracellular taurine levels and impedes mitochondrial respiration in thermogenic adipocytes. Finally, we show that taurine supplementation can alter the chromatin landscape of primary inguinal cells to upregulate genes linked to enhanced NST and metabolic health. Collectively, our study provides novel mechanistic insight into the role of taurine in NST and positions taurine as a potential treatment option to combat obesity and metabolic disease in the future.

## 2. MATERIAL AND METHODS

### 2.1 Animals

Studies on mice were performed according to the permission of the Cornell University Institutional Animal Care and Use Committee. 4-week-old wild-type male C57BL/6J mice were purchased from Jackson Laboratory (#000664) and housed at room temperature (25 °C) with 12-hour cycles of darkness and light. Mice were fed *ad libitum* food and water. To conduct pharmacologically induced thermogenesis experiments, 5-week-old mice were acclimated to a thermoneutral environment (30 °C) for 7 days. Subsequently, mice were daily intraperitoneally injected (IP) for seven consecutive days with either saline or 1 mg/kg of CL 316,243 (Cayman #17499) (n=4, per treatment). Following euthanasia with carbon dioxide, brown, inguinal, and white adipose depots, as well as liver tissue, were collected. Each sample was immediately flash frozen in liquid nitrogen and stored at −80 °C for further protein or RNA extractions.

### 2.2 Cell Culture

Primary inguinal adipose tissue was harvested from 3-week-old male wild-type C57BL/6J mice, followed by thoroughly chopping with scissors for 5 minutes and digesting with 15 ml of lysis buffer (PBS, 1.3 mM CaCl_2_, 2.4 unit/mL dispase II (Sigma D4693), and 1.5 unit/mL collagenase D) in a shaking water bath at 37°C for 15 minutes. Lysates were filtered through a 100 μm cell strainer and spun down for 5 minutes at 600 g at 4°C. After removing the digestion buffer, the stromal vascular fraction (SVF) was resuspended in adipocyte culture media (DMEM/F12 with 10% FBS, 25mM HEPES, and 1% PenStrep) and filtered through a 40 μM cell strainer, followed by centrifuging for 5 minutes at 600 g at 4 °C. Subsequently, cells were resuspended with adipocyte culture media and plated on polystyrene cell culture plates, coated with 2% gelatin. After 48 hours, cells were gently washed with PBS two times and replenished with fresh adipocyte culture media. All primary inguinal and immortalized brown adipose cell cultures were grown at 37°C with 5% CO_2_. For adipocyte differentiation, cells were seeded on polystyrene cell culture plates with 2% gelatin coating. The following day, cells were differentiated with DMEM/F12 (supplemented with 5 ug/mL insulin, 1 μM Rosiglitazone, 1 μM Dexamethasone, 0.5 mM Isobutylmethylxanthine, and 1 nM T3). After 48 hours, the medium was replaced with maintaining media (5 ug/mL Insulin and 1 μM Rosiglitazone) and replenished every two days until Day 6. On the seventh day, cells were treated with 1μM CL 316,243, PBS, or 1mM taurine for different experimental designs.

### 2.3 Generation of Ado-KO cells

The single guide (sgRNAs) of the CRISPR-Cas9-based Ado knockout cells were designed according to the following database https://chopchop.cbu.uib.no/. The sgRNAs sequences are as follows: sgAdo forward: 5’-*TTCCCGGGCCGAGTACACCG*-3’, sgAdo reverse: 5’-*CGGTGTACTCGGCCCGGGAA*-3’. The sgRNAs were cloned into the LENTICRISPR v2.0 (Plasmid #52961) plasmid vector developed by the Zhang’s group. The vector V2 CRISPR DNA Plasmid (1ug) was co-transfected in 293T cells along with 3 µg of the viral envelope PMD2 (Addgene # 12259) and 4 µg of the viral packing PsPAX (Addgene #12260) plasmids using the Polyfect reagent according to the manufacturer’s instructions. The empty vector CRISPR DNA Plasmid was used as a control. After 48 hours, the lentiviral supernatant was collected from the 293T cells and transduced into the immortalized brown fat cell line D.E 2.3 (gift from the Kazak lab). Stable transduced cells were then established via puromycin selection (1ug/mL).

### 2.4 Mitochondria Isolation and Seahorse

The brown fat cell line DE 2.3 cells were differentiated as described above. Cells were then scraped with digestion buffer (300mM sucrose, 5mM HEPES, 1mM EDTA, pH 7.2 with KOH) and lysed with a pre-chilled glass-Teflon homogenizer (13 strokes). Cell lysates were centrifuged at 800 g at 4°C for 10 minutes. The supernatant was then collected and centrifuged for 10 minutes at 8500 g at 4°C to pellet the mitochondria. The supernatant was then removed and the mitochondrial pellet was washed with 1 mL 1x MAS buffer (70mM sucrose, 220mM mannitol, 5mM KH2PO4, 5mM MgCl, 2mM HEPES, 1mM EGTA, pH7.2 with KOH) and centrifuged for a further 10 minutes. The mitochondria pellet was then resuspended with 100 µL MAS buffer, and the protein concentration was quantified by BCA Protein Assay Kit (ThermoFisher #23227). The mitochondrial oxygen consumption rate (OCR) value was assayed by the Agilent Seahorse Bioanalyzer. 10 µg of mitochondria was loaded into XFe24 cell culture plates followed by 20 minutes of spinning at 2000g at 4°C and ultimately refilled 450 µL MAS buffer into each well. The mitochondrial stress test compounds were then administered as follows (final concentration): Port A: Pyruvate (9 mM) and Malic acid (9 mM) (for pyruvate-driven respiration), Port B: Rotenone/Antimycin A (135 μM each). Respirometry data were collected using the Agilent Wave software v2.4.

### 2.5 Protein Extraction and Western Blot

Tissues were lysed with 2% Sodium Dodecyl Sulfate (SDS) supplemented with protease inhibitor (ThermoFisher #A32963) and homogenized with metal beads for 30 minutes at 4°C. Lysates were then centrifuged at max speed for 20 minutes and the protein supernatant was then retained. To extract protein from cells, adipocyte cultures were scraped with 2% SDS lysis buffer and rotated at 4°C for an hour. The cells were then subjected to ultrasonic treatment for 10 minutes with 30 second on/ 30 seconds off (Biorupter) followed by 15 minutes of centrifugation at 4°C at max speed. The protein concentrations of the both the tissue and cell lysates were assessed by BCA Protein Assay Kit (ThermoFisher #23227) and protein samples were supplemented with 4x laemmli blue (Bio-Rad #1610747) and heated at 37 °C for 10 minutes. Prepared samples were resolved on SDS-polyacrylamide gels and then transferred to PVDF membranes. A membrane was blocked with 5% milk for one hour and then washed with TBST before being incubated with primary antibodies overnight. The following day, membranes were washed with TBST and then targeted by secondary goat anti-mouse or anti-rabbit antibodies. Following the secondary staining, the membranes were washed with TBST and imaged by FluorChem system. Densitometry was conducted by the Image J software.

### 2.6 RNA Extraction and Real-Time Quantitative PCR

Tissues were homogenized with Qiagen TissueLyser II in Trizol reagent (Invitrogen), and cells were scrapped with Trizol reagent (Invitrogen). The RNA was extracted according to the manufacturer’s protocol. The concentration and quality of RNA were analyzed using Nanodrop (ThermoFisher) and the reverse transcription reaction was performed by qScript cDNA Synthesis Kit (Quanta Bio). Gene expression analyses was performed by the CFX384 Real-Time PCR System using SYBR Green (BioRad) for the real-time polymerase chain reaction (RT-PCR).

### 2.7 Measurements of Taurine Levels

Differentiated brown fat DE 2.3 cells were washed with 2 mL PBS and processed according to the manufacturer’s protocol (Cell Biolab # MET-5071) to measure the intracellular taurine concentrations.

### 2.8 Metabolomics

For untargeted metabolomics, six-week-old C57BL/6J mice were daily intraperitoneal injection (IP) injected with saline or 1 mg/kg CL 316,243 (Cayman #17499) for consecutive seven days (n=3, per treatment). Mice were euthanized with carbon dioxide, and inguinal tissues were homogenized with 2 mL cold methanol and 4 mL of cold chloroform. Samples were then mixed with 2 mL molecular-grade water. After 5 minutes of incubation, samples were centrifuged at 4°C for 10 min at 3000 g. The polar metabolites were transferred into new tubes and dried under nitrogen flow. Metabolites were resuspended in 30 µL 30% acetonitrile and analyzed by liquid chromatography/ mass spectrometry (LC-MS) on a Vanquish LC coupled with the ID-X MS (Thermofisher Scientific) conducted by the Harvard Center for Mass Spectrometry. Samples or standards were injected into ZIC-pHILIC peek-coated columns (150 mm x 2.1 mm, 5 micron particles, column temperature maintained at 40 °C, SigmaAldrich). The mobile phase was HPLC grade water, 20 mM ammonium carbonate, 0.1% ammonium hydroxide, and 97% acetonitrile. The flow rate at the first 30 seconds ramped from 0.05 to 0.15 mL/min and was maintained at 0.15 mL/min. All data were acquired in the ID-X polarity switching at 120000 resolutions. MS_1_ data was acquired in polarity switching for all samples. MS_2_ and MS_3_ data were acquired by the AquirX DeepScan function for pooled samples. Results were analyzed in Compound Discoverer 3.3 (CD, Thermofisher Scientific). Identifications were based on MS_2_/MS_3_ matching with the mzVault library and corresponded to retention time built with pure standards (Level 1 identification), or on mzCloud match (level 2 identification). For taurine-targeted metabolomics, primary inguinal cells were separately treated with 1 µM CL (Cayman #17499) or PBS for 24 hours. Subsequently, cells and media were processed with the same biphasic extraction described previously. Taurine concentrations were then measured via LC-MS and analyzed the area of the exact mass for the corresponding ions of the targets as well as the intracellular labeled isotopic taurine levels (13C2, 99%; 15N, 98%, Cambridge isotope #CNLM-10253-PK).

### 2.9 Subcellular Fractionation

Differentiated primary inguinal and brown DE2.3 adipocytes were washed with PBS and then scraped with digestion buffer (300mM sucrose, 5mM HEPES, 1mM EDTA, pH 7.2 with KOH) and lysed with a pre-chilled glass-Teflon homogenizer (10 strokes). Cell lysates were centrifuged at 1200 g at 4°C for 10 minutes. The supernatant was then collected and centrifuged for 10 minutes at 8500 g at 4°C to separate the mitochondria. The supernatant was then transferred to a new tube and stored as cytosolic fractions. The mitochondrial pellet was washed with 500 µL mitochondria storage buffer and centrifuged at 4 °C for 10 mins two times (Qproteome Mitochondria Isolation Kit #37612).

Differentiated primary inguinal and brown DE2.3 adipocytes were washed with pre-cold PBS and scrapped with 10 mL PBS. After spinning at 450 g at 4 °C for 5 min, resuspended the nuclear pellet with lysis buffer (Qproteome Nuclear Protein Kit #37582) and incubated on ice for 15 minutes. Subsequently, mixed the sample suspension with 25 μL detergent solution NP (Qproteome Nuclear Protein Kit #37582) and spun at 10,000 g at 4 °C for 5 min. After removing the supernatant, resuspended the pellet with the lysis buffer, followed by centrifuging at 10,000 g at 4 °C for 5 min and storing the nuclear pellet at −80 °C for the protein process.

### 2.10 Nuclei Extraction and ATAC-seq Analysis

2 × 10^6^ primary inguinal cells were seeded in a 10 cm culture plate and scrapped with 1mL PBS after fully differentiating. Subsequently, cells were centrifuged at 4 °C for 10 min at 200 g to generate cell pellets. Cell pellets were then resuspended by 200 µL homogenization buffer (60 mM Tris (pH 7.8), 30 mM CaCl_2_, 18 mM MgAc_2,_ 0.1 mM PMSF, 1mM ß-mercaptoethanol, 320 mM sucrose, 0.1 mM EDTA and 0.1% NP40) and homogenized by plastic pestles for one minute. The samples were then mixed with 1.8mL washing buffer (10 mM Tris-HCl (pH 7.4), 10 mM NaCl, 3mM MgCl_2_ and 0.001% Tween-20) and spun down at 4 °C for 10 min at 500 g. After removing supernatants, the cell pellets were homogenized by the plastic-Teflon in 300 µL washing buffer and filtered through 70 µm nylon filters, followed by spinning at 4 °C for 10 min at 500 g three times. The supernatant was removed, cell pellets were resuspended with transposition mix TD buffer (20 mM Tris-HCl pH 7.6, 10 mM MgCl_2_, and 20 % Dimethyl formamide) and filtered through 40 µm nylon filters. For nuclei quantification, the samples were treated with DAPI (4, 6-diamidino-2-phenylindole) and trypan blue in a 2:1:1 ratio, counted on a hemacytometer. The library preparation and data processing were performed by the laboratory of Dr. Paul Soloway as described previously [30]. The nucleosomal patterning and sequencing was conducted by the Cornell Institute of Biotechnology. The Integrated Genome Viewer 2.13.1 was used to visualize ATAC-seq signal peaks.

### 2.11 Gene Ontology Analysis

Shiny GO v0.741 was used to perform the gene ontology analysis (GO) on the top 50 downregulated and upregulated inguinal gene accessibilities profiles with p-values less than 0.05.

### 2.12 Statistical Analysis

All data were expressed as mean ± SEM, unless other specified. GraphPad Prism 9 was used for the statistical analysis to determine the difference between two independent groups by two-tailed unpaired Student’s t-tests and multiple t-tests. Compound Discoverer 3.2 was used to run the analysis of variance (ANOVA) and Tukey’s HSD post hoc test on the LC-MS data (Thermo Fisher). The significance level was set at p<0.05. Each image legend depicts the value of n as well as the statistical characteristics.

## Supplemental Tables

### 1. Antibody

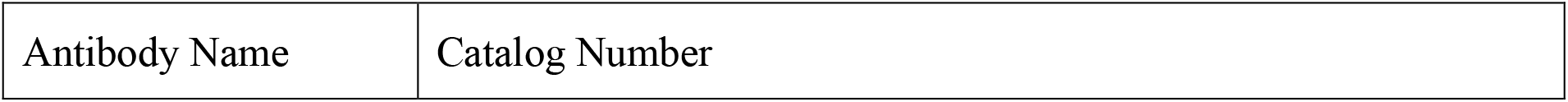
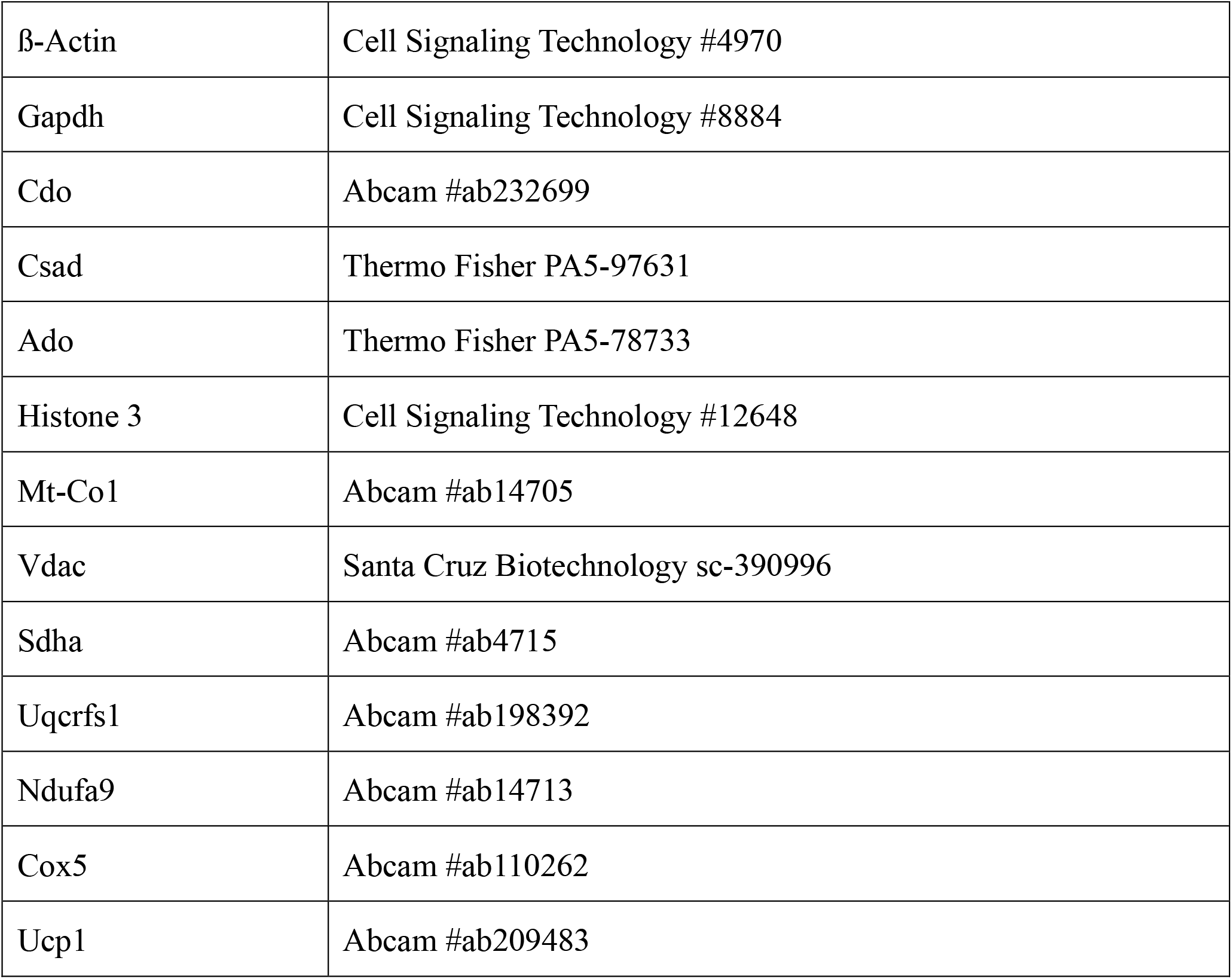

| Antibody Name | Catalog Number |
| --- | --- |
| β-Actin | Cell Signaling Technology #4970 |
| Gapdh | Cell Signaling Technology #8884 |
| Cdo | Abcam #ab232699 |
| Csad | Thermo Fisher PA5-97631 |
| Ado | Thermo Fisher PA5-78733 |
| Histone 3 | Cell Signaling Technology #12648 |
| Mt-Co1 | Abcam #ab14705 |
| Vdac | Santa Cruz Biotechnology sc-390996 |
| Sdha | Abcam #ab4715 |
| Uqcrfs1 | Abcam #ab198392 |
| Ndufa9 | Abcam #ab14713 |
| Cox5 | Abcam #ab110262 |
| Ucp1 | Abcam #ab209483 |

### 2. Primers

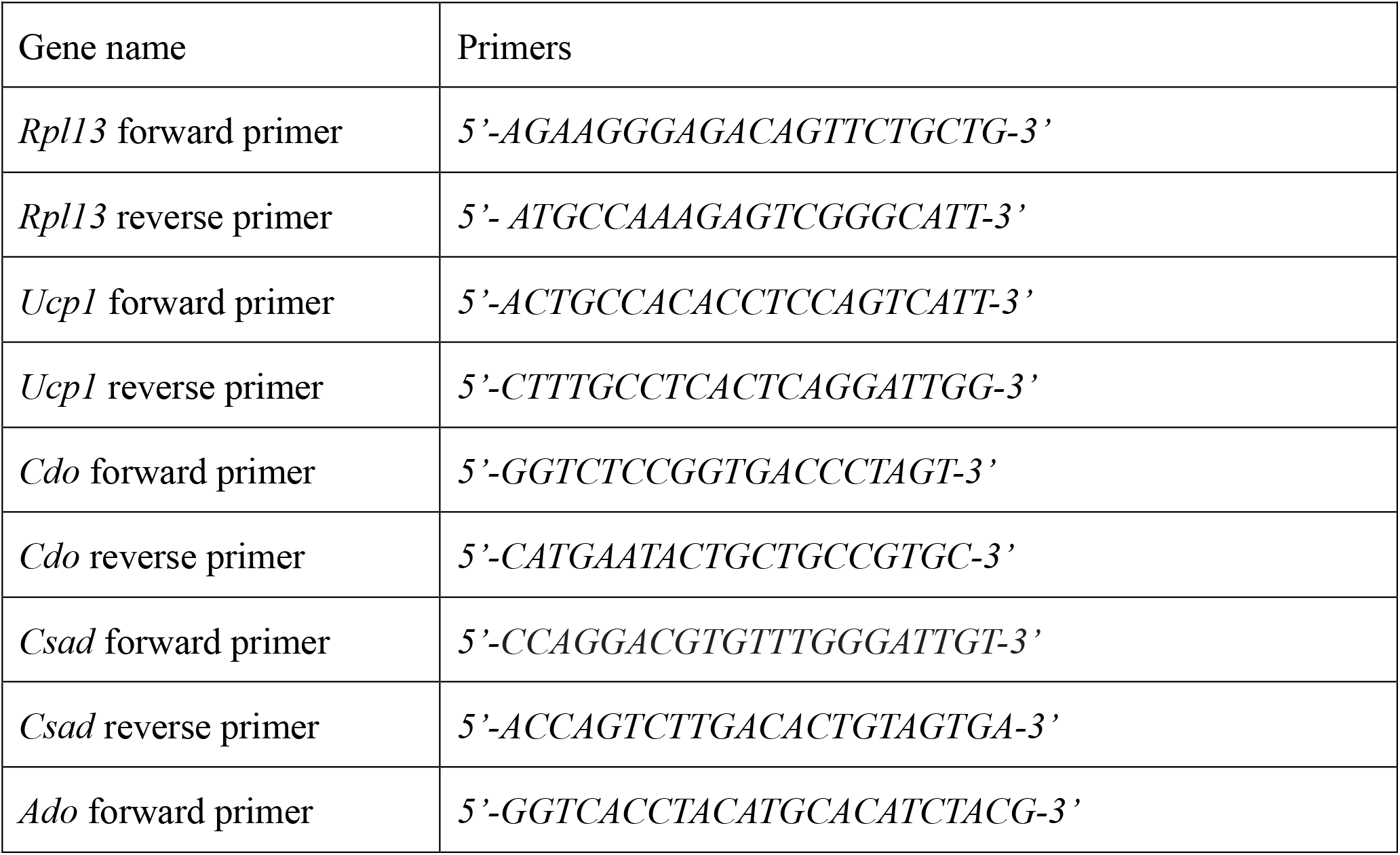
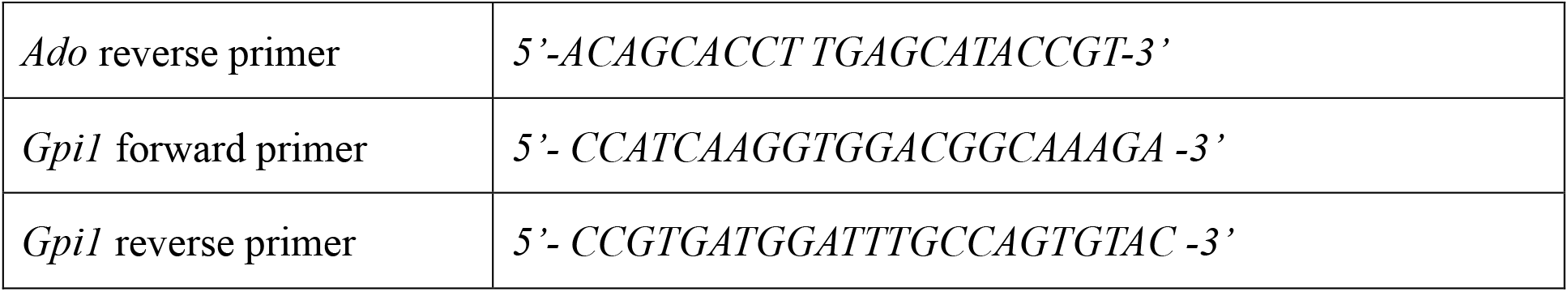

## 3. RESULTS

### 3.1 The taurine biosynthetic enzyme Ado is potently upregulated in inguinal adipose tissue in response to β_3_ adrenergic activation of NST

To investigate the molecular association between taurine biosynthesis and pharmacologically-induced thermogenesis, we injected male C57BL/6J wildtype mice with either the β_3_ adrenergic receptor agonist CL 316, 243, or saline vehicle control for a period of seven consecutive days to activate thermogenesis. We then extracted brown, beige, and white adipose tissue depots in addition to liver tissue and profiled the taurine biosynthetic enzymes to determine their response to activated thermogenesis (Fig. 1A). Taurine can be synthesized from two major pathways. Synthesis can be driven by oxidizing cysteine to cysteinesulfinic acid catalyzed by the enzyme cysteine dioxygenase (CDO) which is then subsequently converted into hypotaurine via the cysteinesulfinic acid decarboxylase (CSAD) to finally generate taurine. Alternatively, taurine can be synthesized from cysteamine via an oxidation reaction catalyzed by the cysteine dioxygenase enzyme (ADO) (Fig. 1B) [31-35]. To determine which taurine biosynthetic enzymes are the most responsive to pharmacologically activated NST, we performed a protein and gene expression profile on the three main adipose depots: brown adipose tissue (BAT), inguinal adipose tissue (IWAT), and white adipose tissue (EWAT) from mice injected with CL. Immunoblot results showed that the protein thermogenic marker Ucp1 increased relative to the saline vehicle control group in all three adipose depots in response to CL confirming that we successfully activated thermogenesis. Protein analysis revealed a significant increase in the protein levels of the taurine biosynthetic enzymes Cdo, Csad, and Ado in the IWAT depots (Fig. 1C). Densitometry analysis further showed that of the three taurine biosynthetic enzymes, Ado was the most robustly induced (Fig. 1C). Interestingly, this was in contrast to the levels of the taurine biosynthetic enzymes in BAT and EWAT which displayed no significant increases after CL stimulation compared to saline vehicle controls (Fig. 1D and E). Gene expression profiling of the IWAT aligned with the protein analysis showing a potent increase in *Ucp1* and all three taurine biosynthetic enzymes *Ado, Cdo, and Csad* (Fig. 1G). Curiously, the *Cdo* and *Csad* as well as the *Csad* and *Ado* mRNA expression levels increased in EWAT and BAT, respectively, but did not align with protein levels (Fig.1H-I). Taurine biosynthetic enzymes are expressed ubiquitously in mammals [35] and to determine if their increase in IWAT is a specific response to NST stimulation or whether it is just a general phenomenon, we interrogated the taurine biosynthetic pathway in the liver. Immunoblot analyses indicated no changes in taurine biosynthetic enzymes in response to NST activation indicating that the robust increases in the gene and protein levels of the taurine biosynthetic enzymes were specific to thermogenic inguinal adipocytes in response to adrenergic stimulation arguing for a specific role of taurine in NST (Fig. 1F and 1J). Taken together, we discovered that the gene and protein levels of the taurine biosynthetic enzymes robustly increase in inguinal adipose tissue in response to β_3_ adrenergic activation of NST. Furthermore, among the taurine biosynthetic enzymes, Ado was the most significantly upregulated suggesting its importance in the taurine biosynthetic pathway.

**Figure 1.**
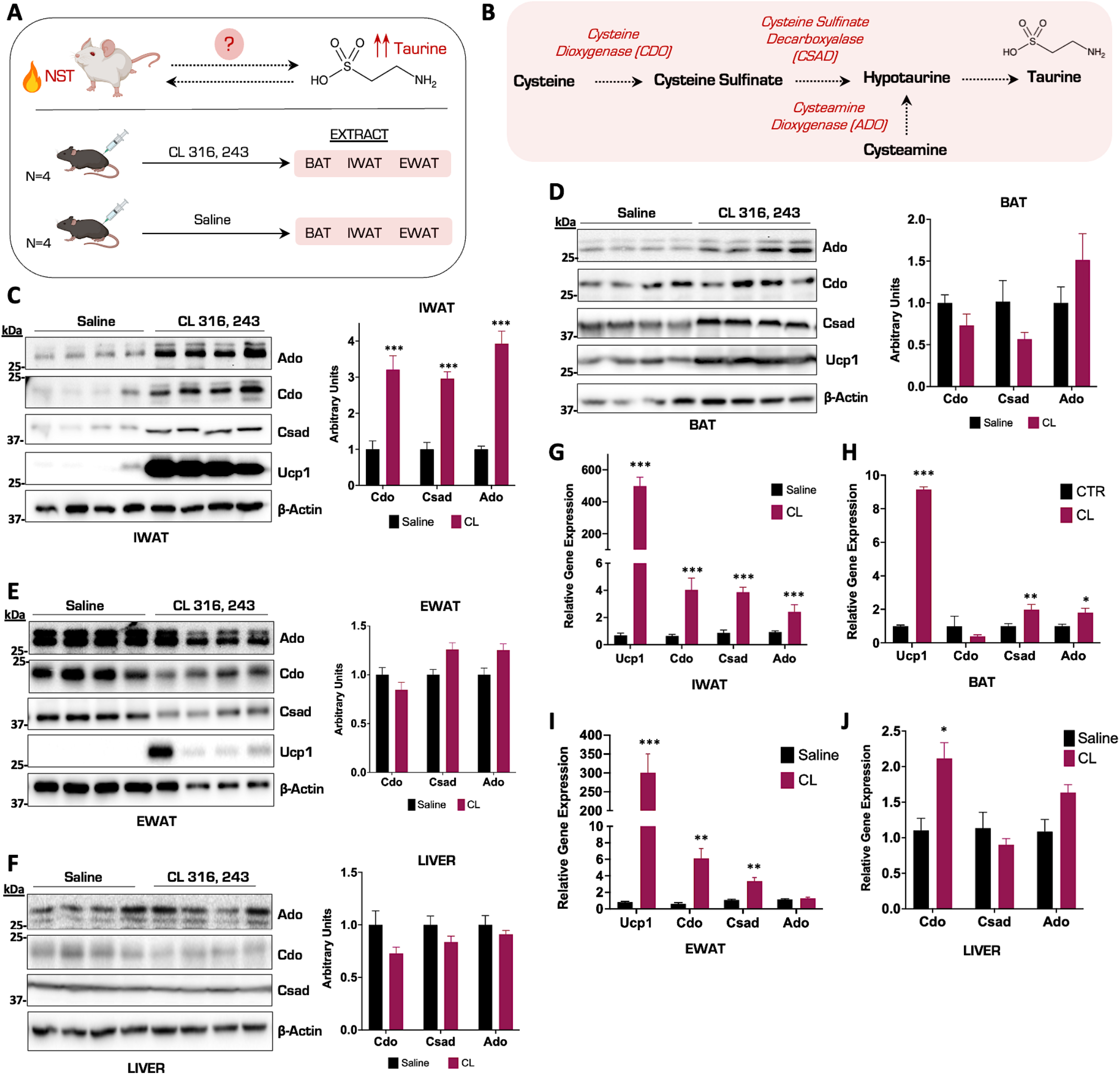
The taurine biosynthetic enzyme Ado is potently upregulated in inguinal adipose tissue in response to β_3_ adrenergic activation of NST. **(A)** Schematic of thermogenesis activation experiment in wild-type (WT) male mice. (N=4) **(B)** Schematic representation of the taurine biosynthetic pathway. **(C-F)** Representative Western blot of IWAT, BAT, EWAT, and Liver tissues from mice injected with CL or saline for 7 days. N=4. β-Actin as the protein loading control. Densitometry of Cdo, Csad, and Ado protein levels normalized to β-Actin are indicated to the right. **(G-J)** Relative mRNA expression of *Ucp1, Cdo, Csad*, and *Ado* from WT mice injected with CL or saline for 7 days in different adipocytes and liver. N= 4. All figures and data are represented as mean ± SEM. Significance is expressed as *p < 0.05, **p < 0.01, ***p < 0.001 by Student’s t-test

### 3.2 Ablation of Ado reduces taurine levels and impairs mitochondrial respiratory capacity in thermogenic adipocytes

Previous studies have shown that taurine levels can be enhanced by NST in IWAT [29] but the magnitude of this increase in relation to cellular metabolites in primary inguinal adipocytes are unknown. We therefore injected mice with CL for seven consecutive days to induce NST and performed untargeted shotgun metabolomics on isolated inguinal tissue to map taurine metabolite levels. Interestingly, compared to energy homeostasis regulators such as nicotinamide adenine dinucleotide phosphate (NADPH) and flavin adenine dinucleotide (FAD), the taurine metabolite levels displayed a significant, albeit modest increase the IWAT (Fig. 2A). We then postulated that perhaps the increase in intracellular taurine levels was modest because the inguinal tissue actively secretes the metabolite into circulation. To test this, we cultured and differentiated primary inguinal adipocytes and stimulated them with CL to activate NST or saline control for 24 hours. We then performed untargeted shotgun metabolomics on both the inguinal adipocytes as well as the extracellular culture media and measured taurine levels. There, we confirmed a significant increase in intracellular taurine levels with no alterations in extracellular media (Fig. 2B and C) indicating that the significant elevation of taurine in response to NST is retained in the cell to perform some unknown function. Given that the Ado enzyme was the most upregulated in response to CL administration (Fig. 1C), we wanted to define the subcellular localization of the enzyme to gain additional mechanistic insight on taurine biosynthesis. We therefore performed subcellular fractionation studies in primary inguinal adipocytes and isolated nuclear, cytoplasmic, and mitochondrial fractions. The Ado enzyme migrates as a doublet in immunoblot analyses yet curiously according to bioinformatic profiling, there is no recorded isoform of the enzyme. Intriguingly, subcellular fractionation experiments revealed that Ado is localized to both the mitochondrial and cytoplasmic compartments yet only the upper band of Ado localized in the cytoplasm while the lower band of Ado is present in both in the cytoplasmic and mitochondrial compartments (Fig. 2D). To define the dependency of Ado on taurine synthesis, we generated CRISPR-Cas9 mediated ablations of Ado in an immortalized thermogenic adipocyte cell line DE 2.3 [36] that displayed the same subcellular localization pattern of Ado as inguinal primary adipocytes (Fig. 2E). We designed two CRISPR single guide RNAs (sgRNAs) termed A1 and A2 targeted to the first exon in Ado (Fig. 2F). The Ado protein and gene levels were fully ablated in the A1 cell population while in the A2 cell line, there was a 50% knock down of Ado with no changes in the other taurine biosynthetic enzymes (Fig. 2G-H). There were also no changes in Ucp1 or the mitochondrial core electron transport chain protein levels as a result of the loss-of-Ado (Fig. 2I). To define the impact of Ado ablation on taurine synthesis, we quantified intracellular taurine levels following a 24-hour CL treatment in the DE 2.3 CRISPR cells compared to vector controls. Complete ablation of Ado (A1) significantly reduced taurine levels while the partial deletion of Ado (A2) did not alter taurine levels indicating that Ado is critical for taurine synthesis (Fig. 2J). We next wanted to assess whether the significant depletion of taurine levels would adversely affect mitochondrial respiratory capacity which is a proxy for functional NST. We therefore isolated mitochondria from both Ado-KO and control DE 2.3 cells and measured mitochondrial respiratory capacity using the Seahorse Bioanalyzer. Mitochondrial respiration was significantly impaired in the A1 cells with complete ablation of Ado compared with control and A2 cell lines (Fig. 2K-L). Overall, we demonstrate that intracellular taurine levels significantly increase in primary inguinal adipocytes in response to pharmacological activation of NST. Furthermore, we show that Ado is localized to both the mitochondrial and cytoplasmic compartments and that CRISPR Cas9-mediated ablation of Ado reduces intracellular taurine levels and correspondingly impairs mitochondrial respiratory capacity, underscoring the importance of taurine in maintaining functional NST.

**Figure 2.**
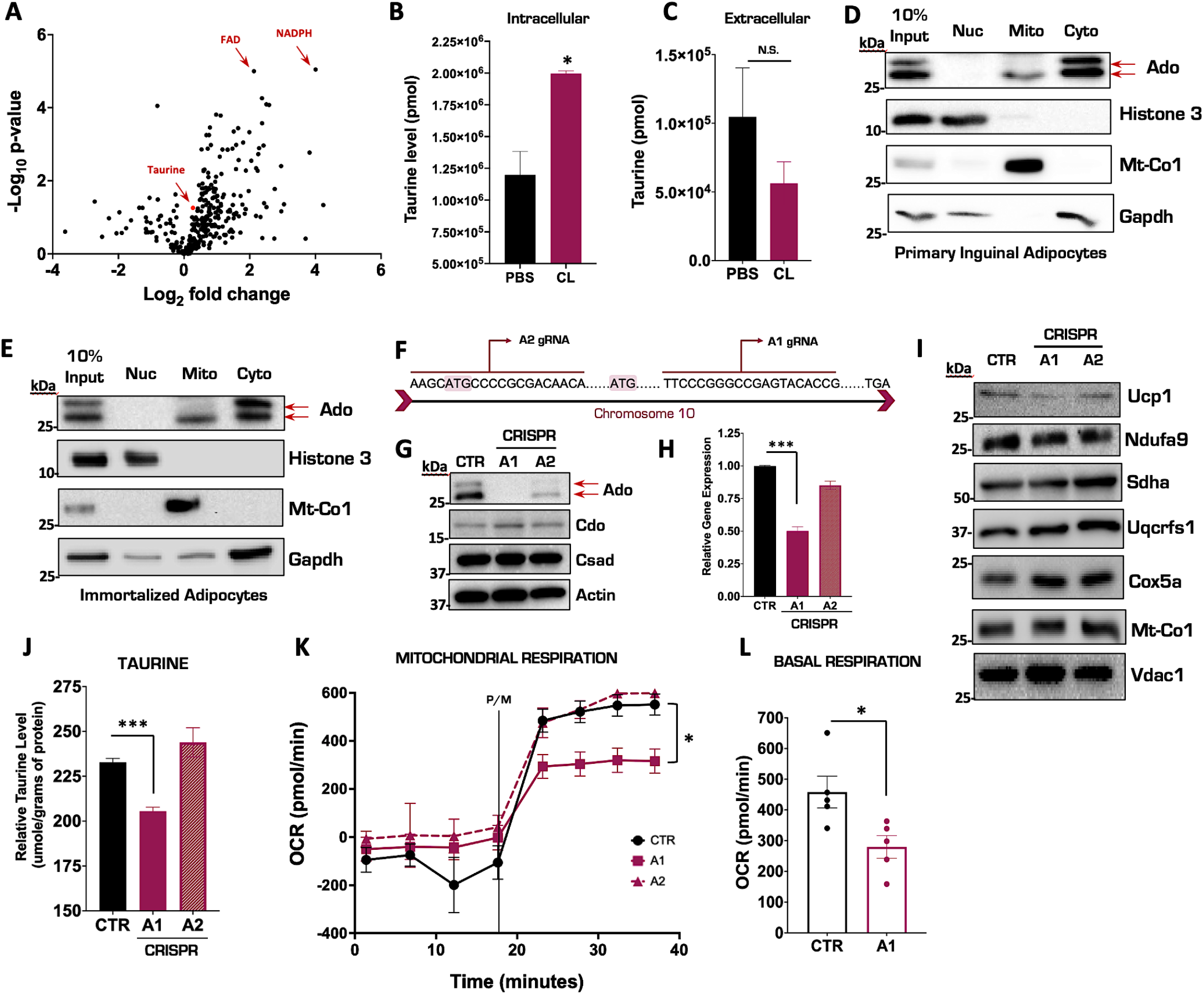
Ablation of Ado reduces taurine levels and impairs mitochondrial respiratory capacity in thermogenic adipocytes. **(A)** Volcano plot of metabolites from IWAT in mice injected with saline or CL for 7 days. The log 2-fold change was calculated based on the ratio of CL versus saline treatment. **(B-C)** Targeted intracellular and extracellular taurine levels in primary inguinal cells treated with CL compared to PBS controls performed by LC-MS. **(D-E)** Representative Western blot for Ado expression in different subcellular fractionations from primary inguinal (D) and immortalized thermogenic DE2.3 cells (E). Histone 3, Glyceraldehyde-3-phosphate dehydrogenase (Gapdh) and Mitochondrially Encoded Cytochrome C Oxidase I (Mt-Co1) represents the nuclear, cytoplasmic, and mitochondrial loading controls respectively. **(F)** Diagram of a portion of the Ado gene sequence and sgRNA binding sites. **(G)** Representative Western blot of Ado CRISPR ablation in DE 2.3 cells. **(H)** Relative mRNA expression of *Ado* in CRISPR DE 2.3 cells. **(I)** Representative Western blot of Ucp1 and differential mitochondrial electron transport chain protein complexes from CRISPR DE 2.3 isolated mitochondria. **(J)** Intracellular taurine levels in CRISPR DE 2.3 cells. **(K)** Oxygen consumption rate (OCR) of CRISPR DE 2.3 vector control and CRISPR Ado isolated mitochondria. (n=5) **(L)** Quantification pyruvate and malate (P/M) induced oxygen consumption rate from CRISPR DE 2.3 control and Ado knock-out isolated mitochondria. The true mitochondrial respiration was calculated by taking the basal OCR values subtracted by the antimycin/rotenone treatment (n=5). All figures and data are represented as mean ± SEM unless otherwise annotated. *p < 0.05, **p < 0.01, ***p < 0.001 by Student’s t test.

### 3.3 Taurine supplementation remodels the chromatin landscape in primary inguinal cells

We have demonstrated that the pharmacological activation of NST results in significant increases in intracellular taurine, but its molecular fate remains unknown. To elucidate a potential mechanistic role for taurine, we hypothesized that taurine may have the ability to alter chromatin structure similar to methionine, another sulfur containing amino acid [37,38]. To test this, we performed an assay for transposase-accessible chromatin with sequencing (ATAC-seq) to map the genome-wide chromatin accessibility in the primary inguinal cells treated with taurine or PBS vehicle control for 24 hours (Fig. 3A). As reported previously, taurine supplementation significantly increases *Ucp1* mRNA levels in primary inguinal adipocytes confirming its ability to successfully activate NST (Fig. 3B). Global ATAC-seq analysis revealed that the taurine supplementation differentially remodeled the chromatin accessibility landscape in several gene loci in primary inguinal adipocytes (Fig. 3C). Gene ontology (GO) analyses of the genes with the most significant upregulated chromatin accessibility patterns highlighted the glucose catabolic and thermogenic metabolic pathways which both play important roles in maintaining NST (Fig. 3D). One of the most upregulated differential gene accessibility patterning was the Glucose-6-phosphate isomerase 1 (Gpi1), which is an enzyme that catalyzes the reversible interconversion of fructose-6-phosphate to glucose-6-phosphate. Taurine supplementation increased the chromatin accessibility of Gpi1 which corresponded with a significant increase in Gpi1 mRNA expression in primary inguinal cells compared to PBS vehicle controls (Fig. 3E and F) [39,40]. This suggests that taurine supplementation can fuel mitochondrial thermogenesis and NST by enhancing glucose metabolism. Overall, we demonstrate that taurine supplementation remodels the inguinal adipocyte chromatin landscape to increase chromatin accessibility and thereby enhancing the transcription of critical genes that promote functional NST. This may be a potential mechanism for the role taurine in NST that can be leveraged towards the treatment of obesity and metabolic disease.

**Figure 3.**
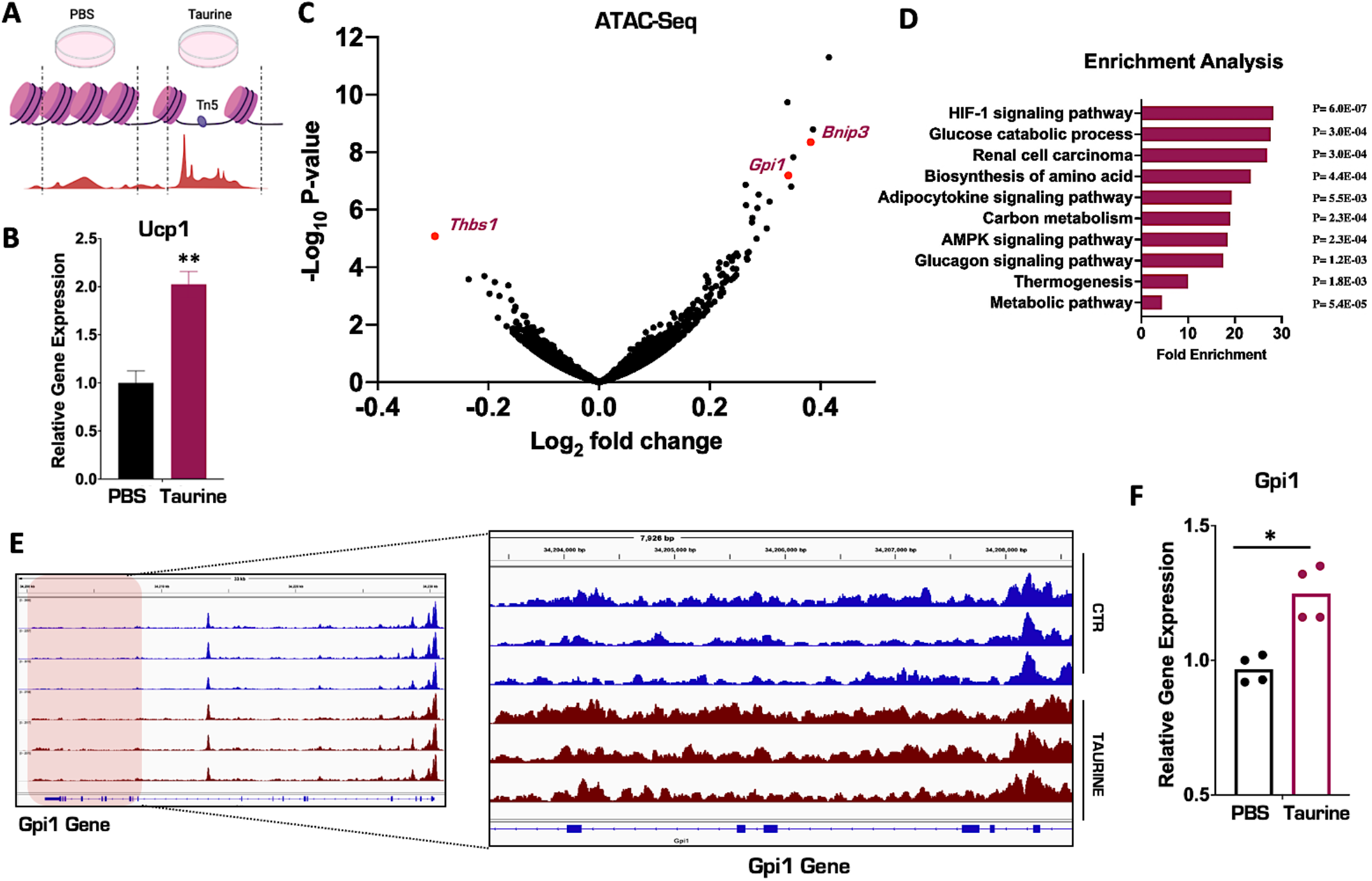
Taurine supplementation remodels the chromatin landscape in primary inguinal cells. **(A)** Schematic of ATAC-seq in primary inguinal cells treated with 1 mM taurine or PBS for 24 hours. **(B)** Relative mRNA expression of *Ucp1* in primary inguinal cells treated with PBS or 1 mM taurine for 24 hours (n=3). **(C)** Volcano plot of differentially expressed gene accessibilities (DEGs) in primary inguinal cells. **(D)** Gene Ontology (GO) Enrichment analysis of DEGs in primary inguinal cells. **(E)** ATAC-seq signal peak calling of the gene Gpi1 in primary inguinal cells treated with PBS (blue) or taurine (purple). To the left: full ATAC-seq chromatin accessibility signal peaks in the Gpi1 gene. To the right: Zoom in of a representative chromatin region (shaded red region) of the Gpi gene to highlight differential chromatin accessibility mediated by taurine treatment. specific chromatin region underlined by the red box. **(F)** Relative mRNA expression of *Gpi1* in primary inguinal cells (n=4). All figures and data are represented as mean ± SEM. *p < 0.05, **p < 0.01, ***p < 0.001 by Student’s t-test.

## 4. DISCUSSION

The integrated nature of taurine and NST remain of great intrigue in the field. Taurine has the ability to activate NST and NST activation correspondingly has the ability increase the biosynthesis of taurine. Indeed, the association between taurine and the protection from obesity and metabolic disease has been well documented [14]. In the current study, we examined the taurine biosynthesis pathway in response to adrenergic activation of NST to determine which adipose depots synthesized taurine. Curiously, of the thermogenic adipose tissues profiled, the taurine biosynthetic enzymes Cdo, Csad, and Ado were only significantly increased in inguinal adipose tissue (IWAT) and not in brown or white adipose depots (Fig 1C). Inguinal adipose tissue also displayed the highest level of taurine upon NST activation. These findings align with those of the field and suggests that taurine may exert its metabolic functions in the inguinal depot.

Taurine biosynthesis can originate from two biological starting points, cysteine-driven synthesis through the Cdo enzyme or cysteamine-driven synthesis through the Ado enzyme. The enzyme Cdo has been well-studied and it has been shown that the loss-of-Cdo in adipocytes significantly reduces taurine levels and impairs mitochondrial function by blunting mitochondrial respiration as well as inhibiting mitochondrial gene expression, such as cytochrome c oxidase subunit 5b (*Cox5b*), ubiquinol-cytochrome c reductase core protein 1 (*Uqcrc1*) and succinate dehydrogenase complex flavoprotein subunit a (*Sdha*) [41,42]. Our study adds to field by providing the first evidence that the biosynthetic enzyme Ado is also critical for taurine synthesis. Ablation of Ado significantly reduces intracellular taurine levels leading to critical failures mitochondrial function and respiratory capacity as mitochondrial oxygen consumption rates were significantly declined (Fig 2J-L). This proves that both the Cdo and Ado taurine biosynthetic pathways are critical to maintain cellular taurine levels. The question however still remains: what is the biological role of taurine in inguinal adipose tissue and how does this metabolite regulate mitochondrial function? It is possible that mitochondrial respiration is impacted due to the defined role of taurine in the modification of mitochondrial tRNAs which is required to translate mitochondrial oxidative phosphorylation (OXPHOS) proteins to generate ATP [43-49]. In our Ado-loss-of-function system however, mitochondrial dysfunction persisted despite the fact that there were no defects with mitochondrial protein translation. Indeed, there were no changes in Mt-Co1 levels in Ado knockout adipocytes compared to vector treated controls. This would suggest that there is an alternative mechanism for taurine in the regulation of mitochondrial function. Another possibility is that loss-of-taurine increased mitochondrial oxidative stress and that this is what impaired mitochondrial function. A previous study observed that depletion of the taurine transporter in murine hearts decreased cellular taurine levels which led to elevated mitochondrial oxidative stress and cellular apoptosis [44]. This is something that will need to be explored in future studies to define the role of taurine in regulating mitochondria function.

In our studies, we noted that the taurine biosynthetic enzyme Ado migrates on immunoblot analyses as a doublet and we confirmed using CRISPR Cas9-mediated ablation that both bands are indeed Ado (Fig. 2G). It is possible that these bands represent different isoforms of Ado despite the fact that alternative isoforms are not reported for the enzyme. It is also possible that there may be post-translational modifications (PTM) on Ado such as phosphorylation or acetylation that may explain the doublet banding. Now that we have established that the Ado-derived synthesis of taurine is critical to maintain taurine metabolite levels in the inguinal depot and is essential to maintain healthy mitochondrial function, the presence of potential PTMs on Ado may shed light into the regulatory roles of this enzyme. This is particularly intriguing since the Ado bands exhibit differential subcellular localization in primary inguinal adipocytes where the upper band is unique to the cytoplasmic fraction while the bottom band is present in both cytoplasmic and mitochondrial compartments [50,51].

It still remains enigmatic why taurine biosynthesis is elevated following either environmental or pharmacological activation of NST (Fig. 2A-B) [12,29]. To our knowledge, taurine is a terminal metabolite and does not participate in any other enzymatic reaction. It also does not become incorporated into protein as the other amino acid building blocks. The function of taurine in inguinal adipocytes therefore remains elusive. In the present study, we postulated that taurine may have an impact on chromatin structure similar to other sulfur-containing metabolites such as methionine. As a DNA methylation donor, methionine can synthesize s-adenosylmethionine (SAM) and induce DNA methylation and correspondingly differential accessibility and expression in specific gene regions [52,53]. We therefore performed a genome-wide ATAC-seq analysis to determine if there are chromatin accessibility changes following taurine supplementation in inguinal adipocytes. Interestingly, we discovered that taurine supplementation can increase the chromatin accessibility of multiple genes associated with NST metabolic pathways such as glucose catabolism. As a representative example, taurine supplementation increased both the accessibility and mRNA transcription of the Gpi1 gene. Gpi1 is a catalytic enzyme involved in glycolysis and multiple studies have proved that both glucose uptake and the glycolysis pathway are enhanced under cold-induced NST in mammals [54-56]. In human studies, PET-scanning also demonstrated that the radio-labelled glucose (^18^FDG) uptake significantly increased in young healthy volunteers, while decreasing in elderly or patients diagnosed with metabolic diseases, who have been considered to have lower NST capacity. Taurine may therefore act as a signal to activate metabolic pathways such as glucose catabolism that would increase the substrates to fuel mitochondrial NST.

In conclusion, we provide evidence to a potential mechanistic role for taurine in the regulation of NST. Following NST activation, taurine levels significantly accumulate in the inguinal adipose depots and its synthesis is driven by all three taurine biosynthetic enzymes of which Ado appears to be the most responsive to NST stimulation. We demonstrate that Ado is critical not only to maintain taurine levels in the cell but that it is also critical for mitochondrial bioenergetic capacity. Finally, we share our findings that taurine supplementation either directly or indirectly alters the chromatin landscape to enhance a suite of genes that promote functional thermogenesis which will lead to protection against obesity and metabolic disease. Taurine can be purchased as an over-the-counter supplement and supplementation is not reported to have adverse effects in humans. Indeed, taurine is currently being tested in several clinical trials including trials to continue to define its efficacy in the protection from obesity [57-61]. Defining the molecular function of taurine will be beneficial to leverage this pathway towards the treatment of obesity and metabolic disease.

## AUTHOR CONTRIBUTIONS

Conceptualization: J.J.B. and P.T.; Methodology: P.T. and J.J.B.; Investigation: P.T., B.S., S.L., Y.L., Y.Q., C.C., K.E., C.W., R.M-K.; Global Analyses: P.T., B.S., S.L., P.S.; Writing: J.J.B., P.T.; Supervision: J.J.B.

## DECLARATION OF COMPETING INTEREST

The authors declare no conflict of interests

## ACKNOWLEDGMENTS

We kindly thank L.K for the immortalized D.E. adipocytes.

## FUNDING SOURCES

Funding for this work was provided by Cornell University Startup Support.

## Notes

### Competing Interest Statement

The authors have declared no competing interest.

